# PhysiBoSS: a multi-scale agent based modelling framework integrating physical dimension and cell signalling

**DOI:** 10.1101/267070

**Authors:** Gaelle Letort, Arnau Montagud, Gautier Stoll, Randy Heiland, Emmanuel Barillot, Paul Macklin, Andrei Zinovyev, Laurence Calzone

## Abstract

Due to the complexity of biological systems, their heterogeneity, and the internal regulation of each cell and its surrounding, mathematical models that take into account cell signalling, cell population behaviour and the extracellular environment are particularly helpful to understand such complex systems. However, very few of these tools, freely available and computationally efficient, are currently available. To fill this gap, we present here our open-source software, PhysiBoSS, which is built on two available software packages that focus on different scales: intracellular signalling using continuous-time markovian Boolean modelling (MaBoSS) and multicellular behaviour using agent-based modelling (PhysiCell).

The multi-scale feature of PhysiBoSS - its agent-based structure and the possibility to integrate any Boolean network to it - provide a flexible and computationally efficient framework to study heterogeneous cell population growth in diverse experimental set-ups. This tool allows one to explore the effect of environmental and genetic alterations of individual cells at the population level, bridging the critical gap from genotype to phenotype. PhysiBoSS thus becomes very useful when studying population response to treatment, mutations effects, cell modes of invasion or isomorphic morphogenesis events.

To illustrate potential use of PhysiBoSS, we studied heterogeneous cell fate decisions in response to TNF treatment in a 2-D cell population and in a tumour cell 3-D spheroid. We explored the effect of different treatment regimes and the behaviour and selection of several resistant mutants. We highlighted the importance of spatial information on the population dynamics by considering the effect of competition for resources like oxygen. PhysiBoSS is freely available on GitHub (https://github.com/gletort/PhysiBoSS), and is distributed open source under the BSD 3-clause license. It is compatible with most Unix systems, and a Docker package (https://hub.docker.com/r/gletort/physiboss/) is provided to ease its deployment in other systems.

## Introduction

Mathematical modelling is widely used to address questions such as tumour progression and tackle the complexity of biological systems [1–3]. Simple and focused models of individual cells can already be very informative to answer a biological question and may be preferable to very detailed models [1, 4] in order to handle cancer complexity and heterogeneity [5]. However, due to the high inter-dependency of the different biological scales driving the development of a tumour, models that explore the interplay between cells and how they relate to the environment are needed to better describe tumourigenesis [5–8]. As a result, multi-scale modelling is being recognized as an important contribution to build a comprehensive mechanistic view of cancer [7, 8].

Several formalisms can be used to model both the individual cell and the population levels (e.g. discrete, continuous, hybrid, etc.) [8–10]. The choice for the appropriate formalism depends on the modelling scope (see for example the summary of different models used in shape homoeostasis in [11]). Some previous modelling efforts have been done with models of cellular automata [12], with the development of Potts models [13, 14], or agent-based models [15, 16], but these approaches have typically focused on the emergent multicellular dynamics after assigning simple, microenvironment-driven single-cell phenotypes, rather than including both the intracellular events and how individual cellular alterations might affect the tumour.

Nevertheless, some attempts have used differential equations to bridge intracellular and population dynamics, but to overcome computational problems, such models need to be kept simple. A model of partial differential equations has been used to explore the transition from one cell cycle phase to another at the population level [17]. Another model uses ordinary differential equations to explore population dynamics [18]. Similarly, other tools have been developed to include Boolean models in a lattice representing the epithelium [19].

Agent-based models are particularly suitable methods for a multi-scale approach, allowing modelers to integrate multiple scales as well as spatial considerations, and providing a mostly intuitive representation of biological systems [20]. Also importantly, these models are very flexible, allowing the simulation of a wide range of situations with minor adaptations. Different multi-scale models have been developed to answer specific questions in the last few years (e.g. [16, 21–24]), but they are usually implemented specifically tailored to a given problem, and they can be difficult to adapt in a straightforward manner to new questions. Additionally, very few of the software packages that allow modelers to combine internal cell signalling with cell mechanics and interactions with its environment are available as open source.

Recently, a promising and open-source software, MecaGen [25], demonstrated the power of combining mechanical behaviour and gene regulation to understand embryogenesis and could be used to study other developmental problems. However, its representation of gene regulatory network as ordinary differential equations (ODE) makes it restricted to small signalling networks. Moreover, due to its morphogenesis scope, MecaGen does not consider cell division dynamics (cell volume growth, cell proliferation, death, etc.), or issues such as clonality (groups of cells within the population with different genetic profiles), which are important in tumorigenesis.

The dynamics study of the cell population is a crucial aspect to improve prognosis or treatment efficiency [6]: knowing the rules governing the behaviour of each separated component in a cell is not enough to predict the emergent behaviour of such a complex system, and similarly the understanding of the processes deregulated in an individual cell is not enough to predict the behaviour of the cell population. The most famous cellular automaton, Conway’s game of life, demonstrates how simple rules that are perfectly known generate unpredictable and complex behaviours at the system level. Moreover, geometry, at the level of a single cell or of the colony, has been shown to have an important effect in the regulation of cell growth [26]. Some studies have demonstrated that genotypically identical cells can adopt different phenotypes according to their environment [27] or tumourigenic factors [28]. Notably, the interplay between spatial position and signalling is critical in development, for example in morphogenesis [29], in cell competition [30] and in cell-fate decision through Notch signalling patterning [31, 32]. Computer modelling is therefore increasingly necessary to tackle such complex problems [33]. The interaction between all these different factors is also crucial to explore the diverse modes of cell motility [34, 35] and is thus a core question in understanding cancer invasion.

Our goal was to develop an open source flexible simulation framework that combines individual cell with intercellular modelling and environment representation, as well as their interaction.

To explore the whole cell population dynamics, the cellular signalling mechanisms and the interplay between cells and their surrounding (i.e., other cells or the microenvironment), we propose to combine an agent-based approach with a Boolean representation of biochemical events taking place in each cell. For that purpose, we have developed a new software, PhysiBoSS, that combines and extends two well-established tools: a signalling pathway modelling tool, MaBoSS [36, 37], which performs stochastic simulations of the signalling pathway inside each cell, is integrated in an agent-based modelling tool, PhysiCell [38], that represents each individual cell as a physical dynamical entity.

We will hereby detail our implementation of PhysiBoSS and present its use with a model of cell-fate decision in response to Tumour Necrosis Factor (TNF) to illustrate the importance of considering cell-cell communication in homogeneous as well as heterogeneous cell population. With this cancer example, we will showcase the use of PhysiBoSS to numerically study the effect of treatment regimes on an heterogeneous cell population and its effects on clonality and tumour growth.

## Design and implementation

To address the issue of including individual cell description into an agent-based model, we adapted, merged and expanded two existing open source software. The first, PhysiCell [38], focuses on the evolution of a tumour by simulating the dynamics of a population of cells under specific constraints, in 2D or in 3D. The second one, MaBoSS [36, 37], defines a contiuous-time Markov process on the state transition graph of a Boolean model and allows the quantification of probability of visiting some selected model states. The two software were merged so that the conditions for the tumour growth depends on the status of individual cells and the behaviour of individual cells is influenced by its environment.

### PhysiCell

The purpose of present work was to simulate not only populations of isolated cells but also organised groups of cells (tissue, spheroid, etc.). Thus, cell-centred, off-lattice models seemed an appropriate choice of agent-based models to be used [9, 39, 40]. Among the available tools implementing cell-centred agent-based models, CellSys [15], Chaste [41] and PhysiCell [38, 42] were particularly interesting.

The implementation of physical laws in CellSys reproduced multi-cellular phenomena quite accurately [43–45]. However, the multi-scale model developed in [45, 46] was restricted to a particular scenario and thus difficult to adapt to other biological questions. Unfortunately, CellSys is not yet distributed as an open-source software that can be easily shared within the community.

Chaste provides an environment to implement different kinds of modelling approaches (agent-based, cellular automaton, vertex model, etc.) within the same framework [41, 47]. Importantly, its implementation allows the direct comparison of outputs from the different modelling techniques and outlines their advantages and limits [40]. However, due to all these possibilities, it is a more complex environment than other agent-based tools. Therefore, to combine it with a network modelling software, we chose to start from an agent-based-dedicated software.

We used PhysiCell source code as a basis for our agent-based model part of our software because of its open-sourced development and its overall simplicity. PhysiCell is a freely available, open-source C++ software (`physicell.mathcancer.org`, [38]). Cells are approximated as spheres which allows for cheaper computation and simpler description than more realistic frameworks. The code is parallelised using OpenMP when possible, allowing the simulation of thousands of cells for several days in a reasonable time (few hours), and simulations of 10^5^ to 10^6^ cells can be run over several days. An efficient implementation of the diffusion of environmental entities (oxygen, glucose, growth factors, etc.) and their interaction with the cells (uptake, secretion, etc.) is also provided by PhysiCell’s BioFVM module [48]. Beyond secretion and uptake of diffusing substrates, PhysiCell has implemeted key phenotypic behaviors needed for our target problems in multicellular systems biology: cell volumetric growth, adhesion, “repulsion”, directed and random motility, cell cycle progression, and apoptotic and necrotic death processes. PhysiCell allows users to attach tailored C++ functions and data structures to each individual cell, which can then modify the cell agents’ phenotypes dynamically throughout a simulation. We use this functionality to add MaBoSS’s signalling model to each individual cell agent, and then to link each agent’s signalling state to its phenotypic behavior.

### MaBoSS

MaBoSS [36, 37] is an open-source C++simulator of Boolean models of signaling pathways. In this logical modelling framework, variables (genes, proteins or specific protein functions) can take two values, 0 or 1, mimicking their activity. Each variable is updated according to the status of its regulating variables, connected by logical connectors AND, OR and NOT. Variable state transition are stochastically calculated from parametrisable rates. One MaBoSS model is thus a Boolean network representation of eventually interlinked signalling pathways. Inputs can be upstream events such as receptor activation and outputs are cell responses to the signalling cascade as cell death, proliferation, migration, etc. The optimized implementation of this formalism allows for computation of a high number of variables in the network (up to 100 variables).

### PhysiBoSS

PhysiBoSS integrates these two software frameworks to obtain a detailed description of each cell’s behaviour and how an alteration in a cell can affect the whole population. There are three main parts in the PhysiBoSS structure:

- BioFVM, a module of PhysiCell software that handles the simulation of one or more diffusing environmental entities [48]. It simulates diffusion, degradation and source of diffusible entities in the extra-cellular matrix (ECM) (Fig 1, green). Space is discretised in a voxel mesh containing information of the local density of a given entity (oxygen, glucose, growth factors, etc.).
- PhysiCell core, that handles the representation of the cells’ mechanics [38] and key phenotypic behaviors. A cell is represented as a sphere with two radii, cellular and nuclear. It can move and interact with neighbouring objects, divide and change its properties according to specific conditions (Fig 1, blue).
- MaBoSS core computes the solutions of a logical model representing the dynamics of a network of intra-cellular events [36]. This module gathers its inputs conditions from the PhysiCell core evaluation (e.g. if a cell has neighbours or if there is presence of growth factors) and retrieves outputs that correspond to cell fates to PhysiCell core (e.g. cell form adhesion, start migrating, dying…). The logical model and parameters description are defined in two files following MaBoSS standard, so any MaBoSS model can be directly used in PhysiBoSS, provided that its inputs and outputs are properly defined and integrated in the agent-based part (Fig 1, orange).

More PhysiBoSS details and specifications can be found in the Supplementary informations (S1 File.

**Fig 1.**
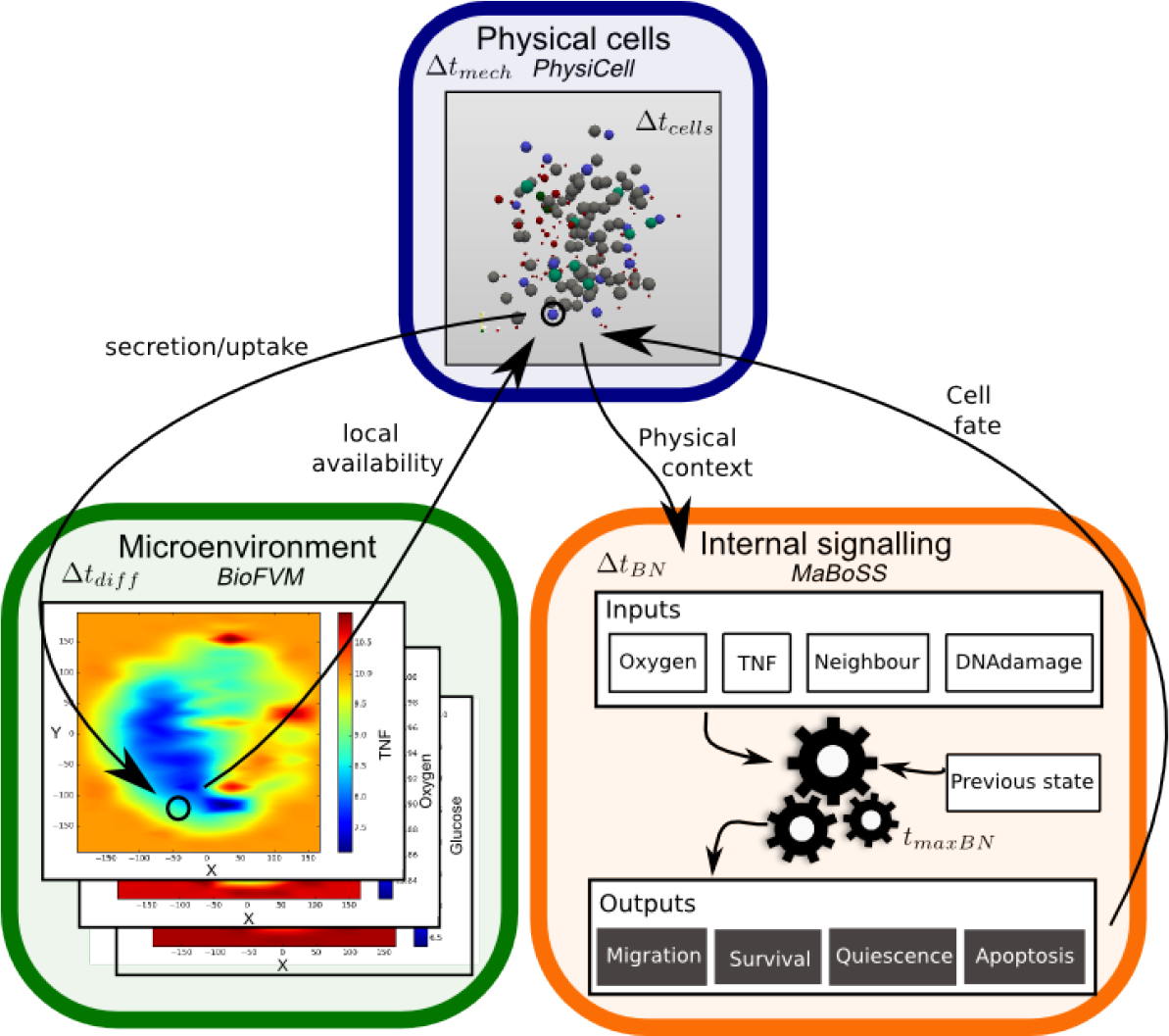
PhysiBoSS structure. Schematic representation of PhysiBoSS: Three main parts are interconnected: the microenvironment representation (green), allowing simulation of diffusing entities; the physical representation of cells as dynamic spheres (blue); and the signalling modelling of each cell (orange).

#### Numerical implementation

The main core of the software is adapted from PhysiCell and the MaBoSS module is compiled as a linked external library. PhysiBoSS is written in C++ with minimal external dependencies (all necessary code is provided within the GitHub repository but the compiler needs OpenMP support). PhysiBoSS is compatible with most Unix operating systems and a Docker container (https://hub.docker.com/r/gletort/physiboss/) is provided to ease its deployment in other systems.

PhysiBoSS uses one executable file that calls an associated case-specific parameter file. This structure is convenient to generate numerous simulations with different parameters/configurations without having to change the code. Moreover, a graphical interface to make the software user-friendly will be developed in the future.

Three executables are provided with present release:

- **PhysiBoSS**, the main executable requires four files: a case-specific parameter file, the two Boolean model files and an initial conditions file. If the simulation is going to take into account extra-cellular matrix, then a matrix-specific initial conditions file is needed.
- **PhysiBoSS CreateInitTxtFile** generates automatically an initial conditions file that specifies initial position of the cells, their volumes and Boolean states for a variety of classic geometries (e.g. sphere, cylinder, rectangle., etc.). For more complex geometries (e.g. Hello World example on PhysiBoSS GitHub documentation https://github.com/gletort/PhysiBoSS/wiki), initial configuration file can be created from binary image of the desired shape by placing cells on the positive areas (script available on GitHub).
- **PhysiBoSS_Plot** generates an `.svg` output snapshot of the simulation at a given time point (more info on GitHub wiki). Options for this plotting are still limited, as the development of visualization and a graphical interface are in scope of future releases of PhysiBoSS.

The preparation, execution and visualization of simulations can be found in detail in S1 File and scripts are provided on the GitHub repository to automate these, along with step-by-step examples with all the necessary files. Time required for one individual run is strongly sensitive to its parameters, such as time/space steps, number of cells, diffusing entities, etc. A Linux cluster to run simulations with OpenMP parallelization was used, but it can also be run locally. Examples of time it required are given in S1 Table.

#### Time scales

Because the system involves a broad range of events at different biological scales, different time scales need to be considered. In particular, reaction-diffusion of biochemical densities in the microenvironment occurs at a very small time scale compared to cell movement, cells gaining volume or cell division (Fig 1). To take these differences into account and avoid unnecessary computation of all scales at all time steps, PhysiBoSS uses PhysiCell’s three time scales (Δ*t*_*diff*_ for diffusion, around 0.01 min, Δ*t*_*mech*_ for cell mechanics, around 0.1 min and Δ*t*_*cells*_ for cell processes, around 6 min; [38]) and adds a fourth one: Δ*t*_*BN*_ (Δ *t*_*BN*_ ≥ Δ*t*_*cells*_, around 10 min) that determines when the intra-cellular Boolean model is updated.

The frequency of the model update Δ*t*_*BN*_ and the length of MaBoSS running time at each update *t*_*maxBN*_ are parameters that need to be carefully studied and set in a case-specific way. As was discussed in MaBoSS paper [36], it is possible to evaluate the network until it reaches one of the stable states, or for a shorter amount of time to take into account transient states. Thus, the MaBoSS evaluation time *t*_*maxBN*_ depends on the biological question of interest and the necessity or not to look at transient events. More details on the interplay of *t*_*maxBN*_ and Δ*t*_*BN*_ can be found in Supplementary informations (S1 File).

#### PhysiBoSS simulations’ features

PhysiBoSS works with spherical **cells** that represent the living cells that can grow/shrink, divide, move, interact with its environment or with other cells, and die. These cells progress through the **cell cycle** and change their physical properties, have a front-rear **cell polarity** and can be part of **cell strains**, where each cell shares a set of common physical and genetic parameters. Again, PhysiBoSS details and specifications can be found in the Supplementary informations (S1 File.

**Simulation of different cell strains** - Using PhysiBoSS parameter file, users can simulate heterogeneous populations of genetically and/or physically different cells. The parameter file must take into account all physical parameters of each strain type, as well as the transition rate of mutated genes of genetically-different strains. PhysiBoSS implements mutation by modifying the variable on-off transition rates, not changing the Boolean network structure. For example, over-expression of a gene will be implemented as a node with very high activation rate and a zero deactivation rate. These transition rate need to be assigned a variable in MaBoSS configuration files and their values need to be specified for each cell strain in the parameter file. Details on how to define all these variants can be found in the GitHub repository together with more examples.

**Extra-cellular matrix representation** - As PhysiBoSS aims to integrate environmental, multi-cellular and intra-cellular descriptions of biology, the representation of the ECM was addressed in present framework. In previous theoretical works, ECM has been represented by a fibrous matrix in a chemomechanical model [49], as cells of a Cellular Potts Model [14, 50], as linear elastic medium [51], as a network of Hookean springs [52], as (non-)deformable objects composed of networks of springs [53], or as passive spheres [15]. The choice for these different representations is strongly dependent on the biological question: a discrete ECM representation can be enough, while in other cases it is necessary to model the deformation, softening, hardening or degradation of the ECM. It is also often a compromise between computational cost and level of precision.

PhysiBoSS proposes two ways of implementing ECM modelling: the first representation is to use ECM as passive spheres (Fig 2A) that can be pushed by other spheres or active cells depending on a friction coefficient. Cells can also degrade (or reinforce) these passive spheres upon contact with user-defined rates reducing (or increasing) their radius. The advantage of this implementation is that it integrates well within PhysiCell code structure and is not in general highly computationally expensive. It can be used, for example, to reproduce the capacity of cells to create tracks in the ECM or to simulate steric hindrance due to Dextran presence in a medium [54]. However, its precision is poor and not very well suited for simulations of filamentous environment, as it’s often the case with ECM. Moreover, if the simulating space is large (or spheres are small), the high amount of necessary passive spheres can drastically increase the computational cost.

**Fig 2.**
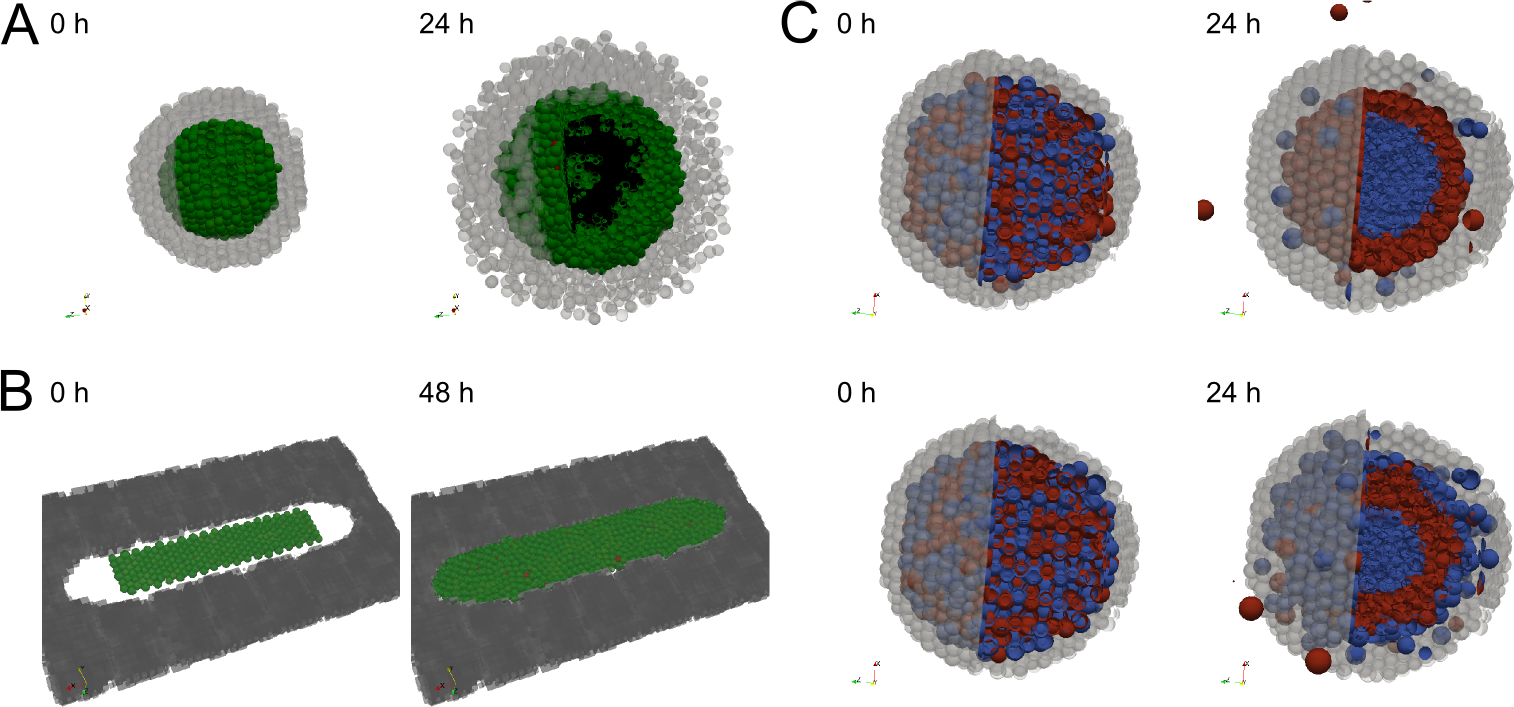
Examples of PhysiBoSS features. A: Snapshots at initial point (t=0) and final point (t=24h) of a simulation of active cells spheroid inside a core of passive cells (grey) with low resistance (can be pushed). B: Snapshots at initial point (t=0) and final point (t=24 h) of a simulation of active cells inside a fixed ECM field (black). C: Snapshots at initial point (t=0) and final point (t=24 h) of a simulation of mechanical cell sorting: the first cell line (blue) forms strong junctions while the other cell line (red) is poorly adhesive, if cells do not adhere to ECM (top) or if the first cell line can attach to the matrix (passive spheres, bottom). A-B: active cells colour are: green, Proliferative cells; red, Apoptotic cells; black, Necrotic cells. A-C: grey: passive spheres representing ECM.

The second representation uses the `BioFVM` module by considering ECM as a not-diffusing density (Fig 2B). Cells can interact with the matrix surrounding them with adherence, repulsion, degradation, deposition of ECM (see details in S1 File), but cannot push it. This allows for a finer spatial ECM definition with small mesh sizes. This representation is very convenient to describe a non-deformable matrix and could be used for example to study cell population growth on restricted areas, as micro-patterns (Fig 2B). However, its non-elastic constraint can be a major drawback for other studies.

**Cell-cell and cell-matrix adhesions** - The core modelling of cell-cell and cell-matrix interactions as presented in [42] are maintained in PhysiBoSS, but with slight modifications allowing dynamic evolution of homotypic, heterotypic [55, 56] and matrix adhesions. Notably, a coefficient of cadherins/integrins density involved in the adhesion was included to respond to the (de-)activation of the Boolean network’s adhesion pathway, so that this coefficient varies accordingly to reflect the different protein recruitment (S1 File).

It has been described that differences in strength between those diverse adhesions can be sufficient to drive specific cell sorting [57–59]. To validate our implementation, we verified that our framework reproduced the sorting behaviour explored in [58]. The results of this can be seen in Fig 2C where the test was limited to a purely mechanical-driven sorting, but having in mind that PhysiBoSS could be used to further explore cell sorting by taking into account cell proliferation and differences in motility, that have been seen to impact the sorting mode or efficiency [60].

## Results

We hereby showcase three examples of scientific problems that PhysiBoSS can address, using a cell fate model upon TNF injection as a case study. This Boolean model started with naive proliferative cells and focused on pathways leading to cellular fates in response to TNF receptor activation, such as survival (read-out of proliferative cells), apoptosis and non-apoptotic cell death (NonACD) [61]. Earlier simulations of this model using MaBoSS predicted that isolated cells in a fixed system lead to heterogeneous fate commitment [61] and this heterogeneity can be interpreted as the limited efficiency of TNF treatment on tumours.

The model used here has been slightly modified to include mRNAs (S1 FigA) of some of the components and adding delays in translation and transcription time-frames [62]. Delays are relevant as they can generate oscillations observed experimentally or differentiation in developing tissue in the case of Notch signalling [63]. Taking advantage of MaBoSS capacity to use time-related parameters, a slower rate on transcription events was imposed and the effects this had on the resulting model behaviour were tested (S2 FigC).

Understanding not only individual cell response but also the dynamics of the collective and environmental interactions is crucial. Thus, we expanded the original set-up from [61] to address important questions on collective behaviours (such as homogeneous or heterogeneous population), spatial (diffusion and consumption of TNF, paracrine secretion of TNF from neighbouring cell, etc.) and dynamical behaviours (continuous or discontinuous presence of TNF, cell proliferation and death, autocrine secretion of TNF through NF*κ*B’s feedback loop, etc.). Moreover, the late appearance of resistant clones within a population justifies the integration of time evolution on the simulation set-up. PhysiBoSS allowed us to address all these questions, drastically increasing agent-based model predictive capabilities.

Different TNF dose regimes in homogeneous cell populations were first simulated using as initial conditions proliferating and healthy functioning cells (S2 File). At frequent intervals, each cell’s internal signalling model was updated according to its current environment (TNF internalization or not) and its current signalling state (resulting from MaBoSS previous iterations). This determined the cell (de-)activation of TNF-α secretion, through NFκB feedback, and the cell fate decision (either to Survival, to Apoptosis or to NonACD, S1 FigB).

In our implementation, cells start as proliferative cells and can evolve to three stable states: Survival (proliferative cells that have activated their NFκB pathway), Apoptosis or NonACD. The internal signalling network of cells committed to Apoptosis or NonACD was not further evaluated, in order to better simulate a dying cell as an irreversible commitment.

### Model validation

To validate our model, two studies focusing on different TNF treatment regimes using 3T3 mouse fibroblast cells in microfluidic chambers were found [64, 65]. These works show that cells’ response to TNF injection is highly heterogeneous. The proportion of cells that responded to TNF injection within the first 8 hours on average (reported in their experiments as transient relocation of NFκB to the nucleus) depended on the dose concentration and the duration of the injection, referred by the authors as ‘stimulus area’ (figure 1b of [64]). The fraction of responding cells varied from 0, for a dose area smaller than 10^2^ ng.s/mL, to a total response when dose area was around 10^4^ ng.s/mL, with a Hill-like dependency on the stimulus area: Hill coefficient being around 1.5 and with a response time from 20 to 50 min [65].

We first calibrated our model to simulate growth dynamics of 3T3 cells. In the absence of TNF, the population grows without constraints (Fig 3A), with a doubling time of approximatively 16 hours [66]. We then explored the response of the population when TNF was injected in the medium (refer to S2 File for details on TNF dynamics). Upon limited TNF injection, only a partial response of the population was observed, in accordance with experimental observations (Fig 3B). We then varied both the TNF concentrations and the injection durations and obtained a similar Hill-like dependency of the fraction of active cells to injection area (Fig 3C). Parameters of TNF dynamics were chosen to have similar range of response to similar injections’ doses of experimental data: no response under 10^2^ ng.s/mL to a “total” response above 10^3^ ng.s/mL, Hill coefficient of 4.8 and a response time between 40 to 60 min.

We measured the activated fraction of cells present at the end of the simulation (i.e. when NFκB got activated at least transiently) as done in the experiments. However, in our simulations, 20%of cells internalized TNF and committed to Apoptosis without activating NFκB pathway. In fact, authors showed that after 8 hours of high TNF concentration (10 ng/mL) cells started to express less anti-apoptotic genes and more pro-apoptotic genes (figure 2a of [64]).

The simulated response was stiffer than the experimental one, but reproduced qualitatively the observed behaviour within the same range of values. This suggests that our model can be used to predict qualitatively the cell population response to TNF in other conditions and that the simulations can retrieve a range of TNF concentration values and percentage of cells that responded, but researchers should be cautious using these as exact values.

### Multicellular spheroid response to TNF treatment

*In vitro* multi-cell spheroid models are now widely used in tumourigenesis [8, 67], due to their similarity with *in vivo* conditions allowing for a good compromise between system complexity and clinical relevance [68]. Therefore, our model was used to investigate how a multi-cell spheroid would respond to TNF injection, and showed that it could be used to test the effect of injection frequency, clonality or complex heterogeneous scenarios.

In the absence of TNF, the spheroid grew as cells doubled their volumes and divided (Fig 4A). In this simulation, the focus was only on the TNF effect and oxygen and nutrient diffusion were not taken into account, which could limit the spheroid growth [69], as we will see in next section.

Continuous injection of a low dose of TNF drastically reduced the expansion of the population (from 4.5 fold increase in cell numbers after 24h in the non-treated simulation to only 1.8 in the treated one) as around 50 % of the initial population committed to Apoptosis or NonACD in response to TNF (Fig4B). However, cells that activated the survival NFκB pathway became resistant to TNF (Survival stable state) and transmitted this resistance to the daughter cells, as these inherit their mother cell’s signalling network state. This sub-population continued to grow independently of the TNF presence: discontinuing the TNF injection or increasing 10-fold the TNF concentration after 600 min did not affect the overall behaviour (Fig 4C). In the first 600 min, these cells received a constant external input (TNF activation) and reached a stable state (as could be predicted from a MaBoSS simulation of an individual cell [61]). In the first scenario, increasing the TNF dose did not affect the signalling network of these already activated cells or their stable state. In the second scenario proliferative cells were still present as, due to the absence of TNF, cells switched from a NFκB pathway-activated proliferative stable state to an un-activated proliferative stable state.

**Fig 3.**
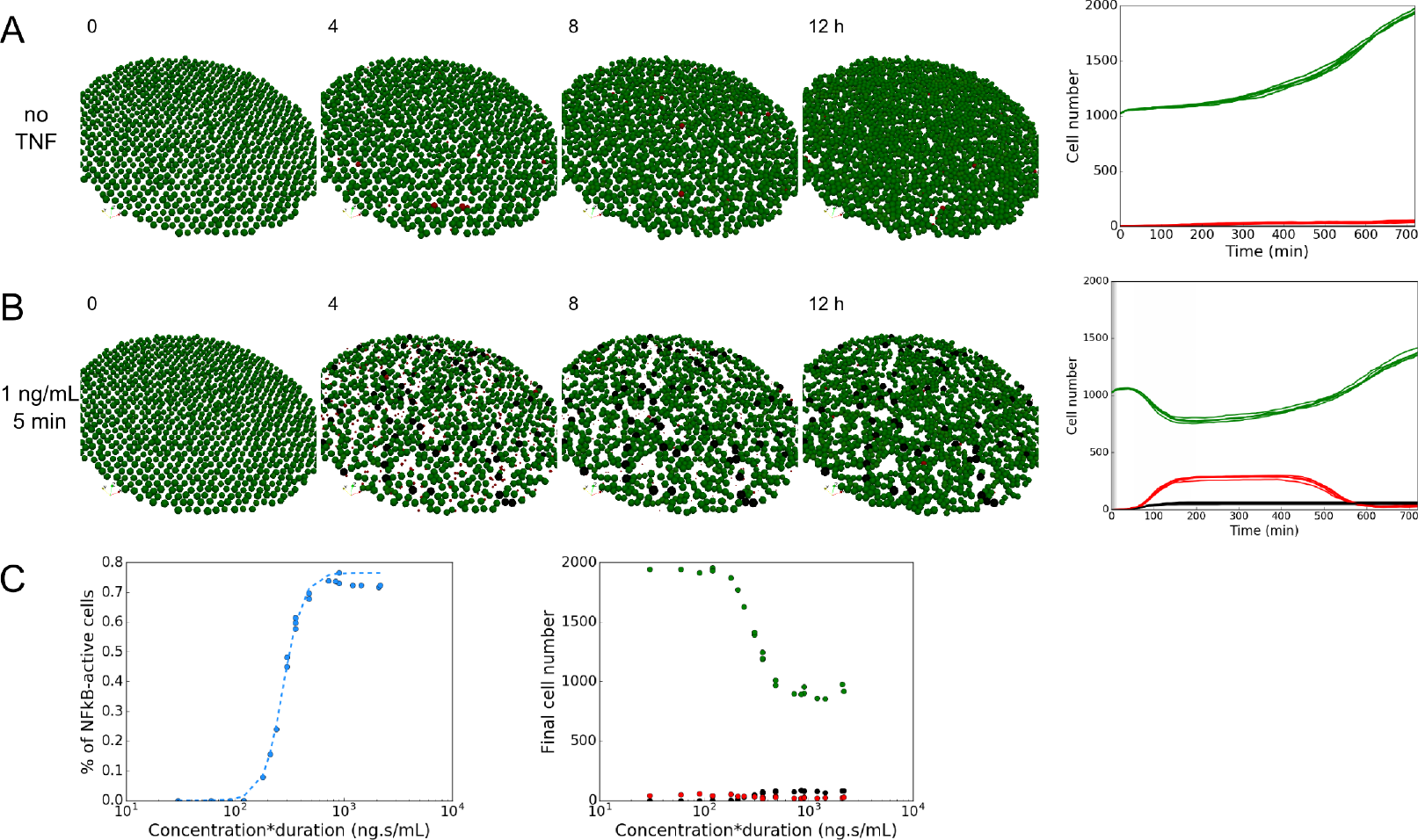
Population response to TNF injection. A: Simulation without TNF. Snapshots of a simulation (left) at t = 0, 4, 8, and 12h. Time evolution of the number of cells in each cell fate (right) for 5 simulations. B: Simulation for a low-dose injection of TNF (1 ng/mL during 5 min). Snapshots of a simulation (left) at t = 0, 4, 8, and 12 h. Evolution of the number of cells in each cell fate during time (right) for 5 simulations. Grey shading represents the time of TNF injection (right), in this case 5 min starting at t = 0. C: Cell population response to TNF stimulus area (concentration time duration of injection). Fraction of “activated” cells (transient NFκB activation) compared to the initial number of cells according to TNF stimulus area (left). The dotted line represents the Hill-function fit (coefficient 4.8). Final number of cell in each fate according to the stimulus area (right). A-C: Green, Proliferative cells; Red, cells committed to Apoptosis; Black, cells committed to NonACD. Simulation time is 12h, initial disk radius is 400 μm, which accounts for roughly 1000 cells.

From these results, we hypothesized that varying the injections of TNF instead of having a continuous regime could sensitise the cells to TNF. To test that idea, we simulated pulses of TNF injections at given frequencies and found out that transient exposure did affect strongly the population’s response (Fig 4D). This is important to consider as *in-vivo* tissue cells are subjected to bursts of TNF expression from neighbouring immune response cells, while they might be under continuous injection in *in-vitro* trials. Cells that were proliferative after the first injection were still responding to TNF in the following injections and the proportion of dying cells was much higher than with a continuous treatment, showing the importance of considering non-steady regimes. Accordingly, the difference between transient and continuous exposures had been observed in a recent in vitro study [70]. This suggests that PhysiBoSS can be used to screen frequencies and concentrations of treatment injections, narrowing *in vitro* investigations.

Notably, cells activated apoptotic pathway in response to the first injection whereas later injections committed cells mostly to NonACD (Fig 4D), which highlighted the importance of dynamics in cell’s response. This was consistent with the construction of our network, with faster apoptosis commitment (S2 File. Moreover, changing the transcription rate in the model affected the type of cell fate decision, as increasing it favoured necrosis (NonACD) (S2 FigC). Indeed, faster transcription caused faster mXIAP activation that caused Apoptosis inhibition and benefited NonACD cell fate (S2 File).

**Fig 4.**
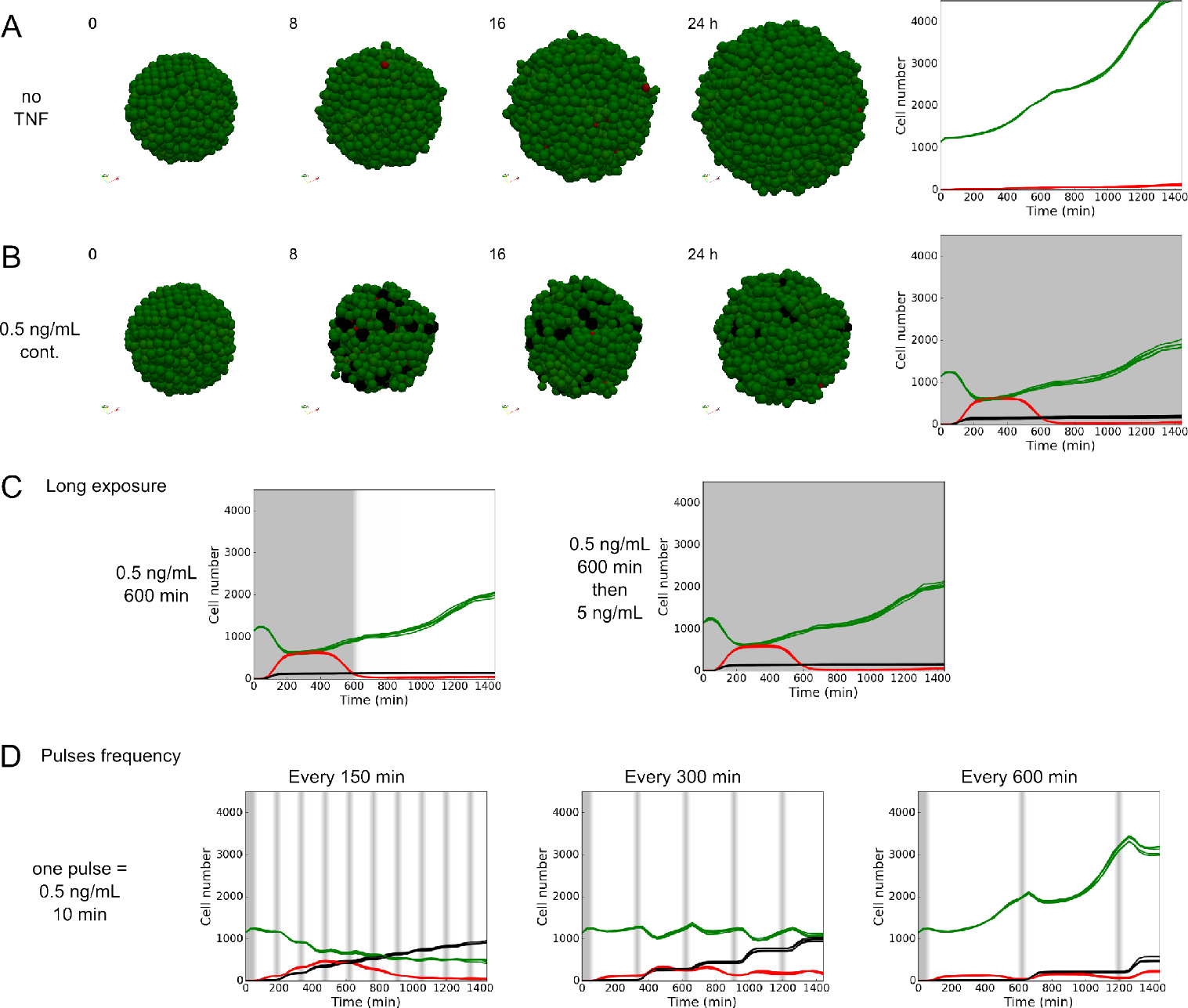
Spheroid response to TNF injection. A: Simulation in the spheroid model without TNF. Snapshots of a simulation (left) at t = 0, 8, 16, and 24h. Time evolution of the number of cells in each cell fate (right) for 5 simulations. B: Cell fates, simulation for a continuous low-dose injection of TNF (0.5 ng/mL, continuously). Snapshots of a simulation (left) at 0, 8, 16, and 24h. Time evolution of the number of cells in each cell fate during time (right) for 5 simulations. C: Simulation when TNF injection (0.5 ng/mL) is discontinued (left) or drastically increased (5 ng/mL, right) after 600 min. Time evolution of the number of cells in each cell fate for 5 simulations under each condition. D: Effect of pulse injection frequencies in the model simulations. Time evolution of the number of cells in each cell fate for 5 simulations when pulsed injections (0.5 ng/mL during 10 min) are repeated every 150 (left), 300 (middle) and 600 (right) min. A-D: Green, Proliferative cells; Red, Apoptosis; Black, NonACD. Grey shading represents the time of TNF injection. Initial spheroid radius is 100 μm, roughly accounting for 1100 cells.

Furthermore, PhysiBoSS can be used to test the system’s response in different cell types, that have different TNF secretion rates in response to NFκB activation, which would affect the overall sensitivity to TNF concentrations (S2 FigA). This tool could also be used to test the effect of the initial size of the spheroid on the TNF availability for cells, and how this would affect the global behaviour, although initial tests using different ranges did not yield different results (S2 FigB).

### Response to TNF treatment of heterogeneous multi-cellular spheroids

One major challenge in tumour treatment is the high level of heterogeneity, spatial and temporal [71], within the population, notably the presence of different clones, which will respond differently to the same conditions. To illustrate the effect of treatment on a genetically heterogeneous population, we simulated a spheroid initially composed of 75% of wild type strain (non mutated, WT) and 25% of mutated cells with IKK and cFLIP over-expressed(+) (Fig 5A). This double mutation was found to drastically promote cell survival (Fig 5B and S2 File) using our pipeline on computational tools for logical models exploration [72]. As expected, part of the WT population died under TNF treatment while the mutant population survived and proliferated. Importantly, the presence of the mutated population did not impact the response of the WT population: the final ratio of surviving WT cells compared to their initial number was similar to the one in a WT-only population (S3 FigC, no significant difference under Kolmogorov-Smirnov test). This observation was also valid with two other mutations promoting either apoptosis or NonACD (S3 FigA-C), or with different initial proportion of WT cells in the total population (S3 FigD,E).

**Fig 5.**
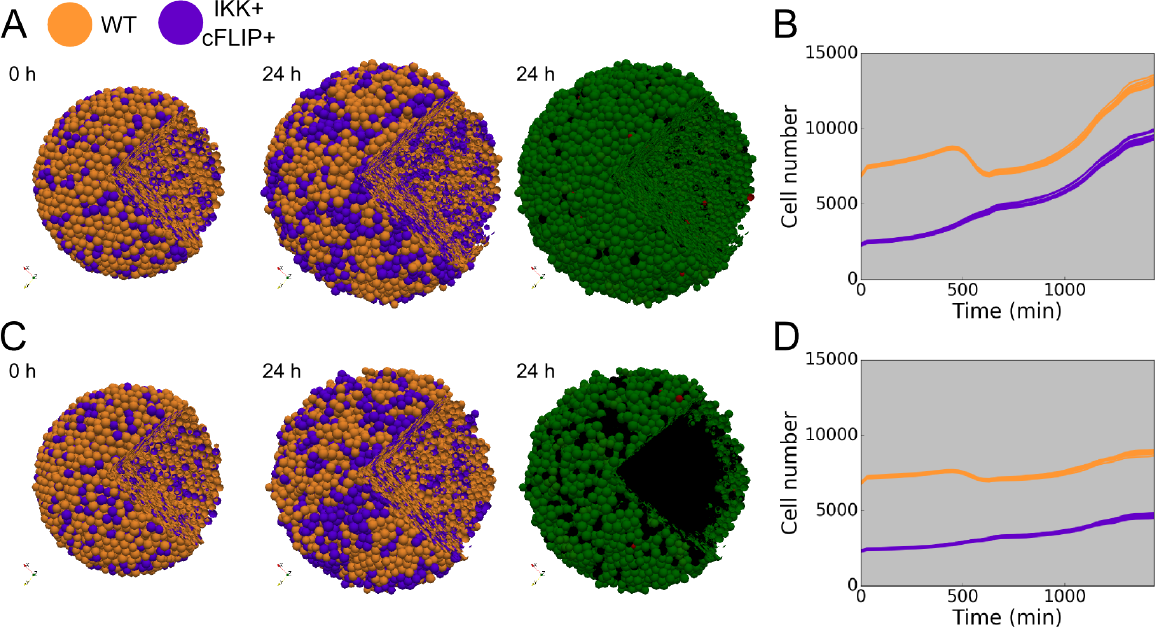
Genetically heterogeneous population under TNF treatment. Simulations of heterogeneous population composed of 75% of WT cells (orange) and 25% of IKK+ and cFLIP+ mutated cells (purple). A: Snapshots of a genetically heterogeneous population simulation at initial and final time (24 h), with cells coloured by cell type (left and middle) or by cell fate (right). B: Time evolution of the number of cells in each strain (WT and mutated) for 10 simulations. Grey shading indicates presence of TNF in continuous injection at 0.5 ng/mL. C: Same as A with oxygen dynamics taken into account. D: Same as B for simulations with oxygen diffusion. A-D: Cell fate colours: green, Proliferative cells; red, Apoptotic; black, NonACD. Initial spheroid radius is 200 *μ*m, which accounts for roughly 9000 cells, + stands for over-expression.

In this simulation, cell communication was limited to TNF consumption/secretion and physical interaction. However, it is known that in crowded environment such as tumours, there is cell competition for resources, e.g. oxygen, nutrients or growth factors. To study the impact of resource competition among cell strains, we included oxygen diffusion and its cell consumption in the 3D spheroid set-up. A threshold under which cell commits to necrosis (NonACD) due to lack of oxygen was fixed (S2 File, Parameter table). As a consequence, in an homogeneous WT population without TNF, a necrotic core formed with a thin proliferative rim around it (S4 FigA), as described for large spheroids [67, 73]. Under TNF treatment, the homogeneous population growth was strongly limited by the combination of TNF-and oxygen-mediated death (S4 FigB).

When mixed with the IKK+ and cFLIP+ mutated population, WT cells had to compete with proliferating mutants cells apart to having to survive to TNF signalling. Thus, majority of the WT cells that did not commit to Apoptosis or NonACD by TNF signal, committed to NonACD by lack of access to oxygen (Fig 5C-D). WT strain growth was considerably reduced compared to its growth in the homogeneous spheroid (S4 FigE). Similarly, when mixed with pro-apoptotic or pro-NonACD mutants, WT cells gained access to oxygen and proliferated more than in an homogeneous spheroid (S4 FigC-E). This competition among strain was still valid for other ratio of WT/mutated population tested (S4 FigF).

The resulting tumour is the consequence of the different adaptive abilities of these strains to the environmental conditions and to the TNF treatment. As illustrated here, this may result in the growth of resistant clones that gain access to nutrients. Importantly, it was recently shown that the tumour spatial structure strongly impacts the adaptive therapy efficiency [74]. Hence, use of tools such as PhysiBoSS that include both spatial distribution and signalling networks are necessary to explore and predict the best clinical adaptive therapy strategies [75].

### PhysiBoSS performs genotype-to-phenotype simulations

PhysiBoSS bridges gene perturbations to cell population dynamics taking into account environmental perturbations. It is thus a unique tool to address issues as clonality in tumours, taking into account both intra-clonal (through stochasticity within MaBoSS and PhysiCell) and inter-clonal heterogeneity, their interaction with the environment (TNF, oxygen, etc.), and their temporal evolution [71]. In particular, PhysiBoSS allowed us to showcase a strong difference of the population’s response to perturbations in TNF availability, which showed to be dependent on the signalling pathway dynamics (see also [70]). Our results also illustrated differences between experimental and *in silico* 2D (≈ microfluidic system) and 3D (≈ spheroid culture) systems, due to differences in access to chemicals, such as oxygen, nutrients or injected drugs. This suggests that conclusions drawn in a 2D set-up could be different from the ones drawn in a 3D environment.

Multi-scale models have a great potential to study morphogenetic events [25, 76] by combining biochemical patterning with cell signalling and mechanics. Indeed, the overall population organisation can be influenced both by cell differentiating in response to external signals (e.g. growth factor access) and cell organisation by mechanical clues (e.g. differences in adhesion or motility). Computational tools such as PhysiBoSS can be used to predict the resulting organisation of such interplay between genetic and phenotype factors under environmental perturbations, and thus reduce experimental exploration [33].

## Availability and Future Directions

PhysiBoSS is available on GitHub (https://github.com/gletort/PhysiBoSS), under the BSD 3-clause license. A Docker image has been created to allow users to run PhysiBoSS even if their system is not compatible with it. We provide in the repository the source code of PhysiBoSS, scripts to use it and analyse results, and extended documentation on its Wiki for installation, usage, and step-by-step examples. We provided in particular detailed examples (with all necessary files) to simulate cell sorting by differential adhesion, spheroid growth under TNF treatment, cell population growth under TNF treatment for an customized initial state (here in shape of Hello World), cells embedded in an ECM field and a cell population composed of three different mutants.

One limitation of PhysiBoSS is its representation of cell constraint to a spherical shape. In the next version of PhysiBoSS, we plan to propose an ellipsoidal shape (for more or less elongated cells), as presented in two recent multi-scale models [25, 76]. We also plan to extend the representation of the extracellular matrix, so that users could choose different modes of implementation according to the biological questions. Indeed, PhysiBoSS will be modified so as to offer different levels of representations from a very abstract representation (as currently possible as a field or passive spheres), to a more realistic representation (e.g. filamentous environment, which could be done by introducing a finite element mesh for the ECM, as suggested in [38]).

Some further extensions of PhysiBoSS will include: (1) the development of the interface between the agent-based part and the Boolean network by adding more possible input or output (IO) nodes (S1 File); (2) the creation of binary output files (instead of txt) with an executable that read, plot and analyse the files within PhysiBoSS; (3) some modifications of MaBoSS implementation to simulate multiple instances at the same time and thus have the possibility to run the network updates in parallel; and (4) a graphical interface to launch simulations in a more user-friendly way (it is currently developed for command line usage only).

Finally, another direction could be to combine MaBoSS with other agent-based modelling software, to allow for choice between different frameworks (e.g. Cellular Potts, Vertex Model) according to the biological question of interest.

Any contributions to PhysiBoSS development are welcomed.

## Acknowledgements

This work received funding from the European Union Horizon 2020 research and innovation program under grant agreement No 668858 (PrECISE project). This work has also been funded by INVADE grant from ITMO Cancer (Call Systems Biology 2012).

## Authors contributions

Conceptualization: GL, AM, PM, AZ, LC; Methodology: GL, AM, LC; Software: GL; Software testing: GL, AM, RH, LC; Visualization: GL; Funding acquisition: AZ, EB, LC; Investigation: GL, AM, GS, LC; Writing - original draft: GL, AM, LC; Writing - review & editing: GL, AM, GS, RH, EB, PM, AZ, LC; Project administration: EB; Supervision: LC.

## Supporting Information

**S1 Fig.**
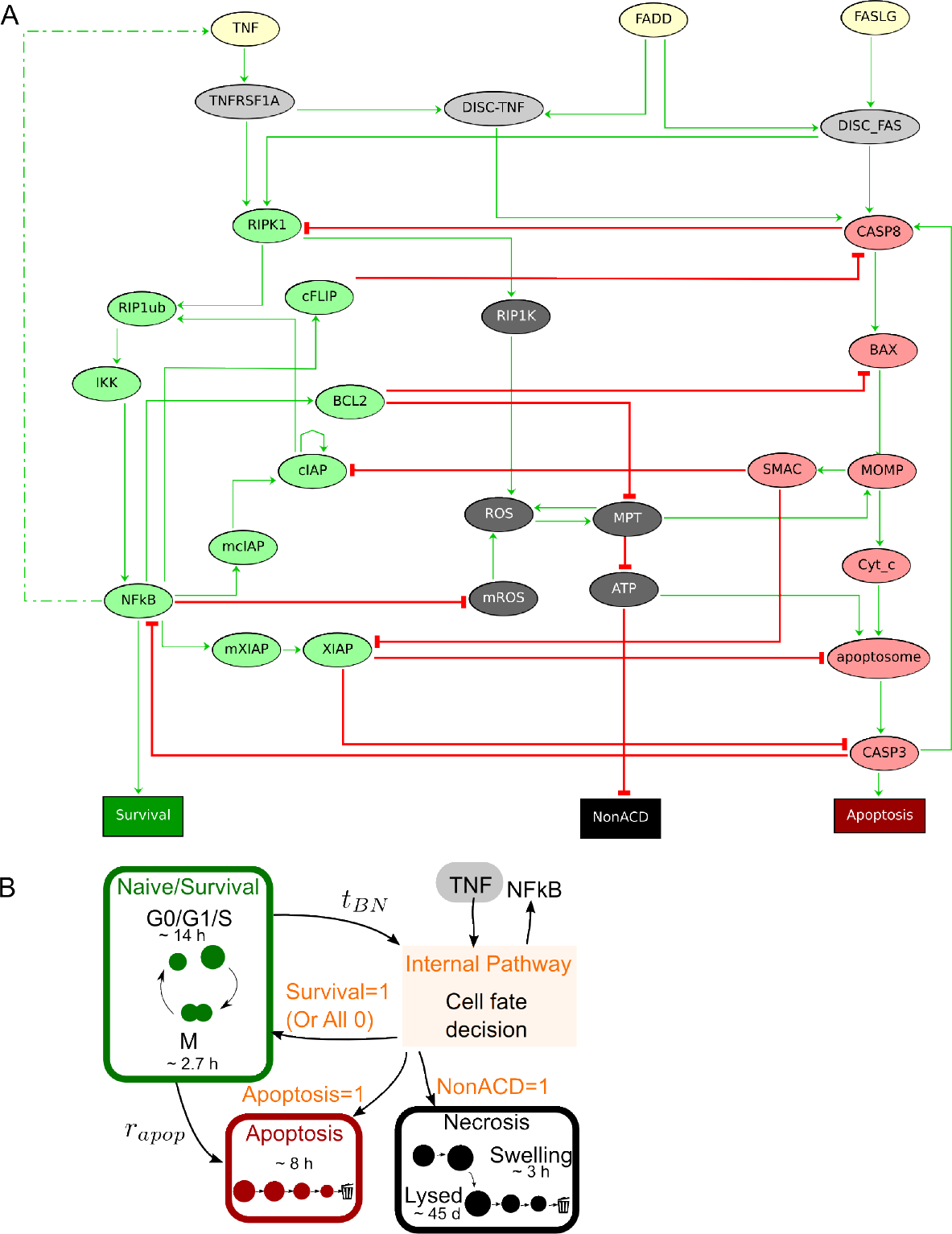
A: Boolean network used. Pathway related to Survival (green), related to Apoptosis (red) and related to NonACD (black). Green arrows represent activation, red arrows inhibitions. B: Schematic representation of how cell cycle is simulated. Cells are initially in Proliferative state (growing and dividing; green). At frequent interval, their internal signalling network is updated by running MaBoSS (orange) given its environmental conditions (internalization of TNF). This decides of the fate of the cell, to continue proliferating or commit to Apoptosis (red) or NonACD (black).

**S2 Fig.**
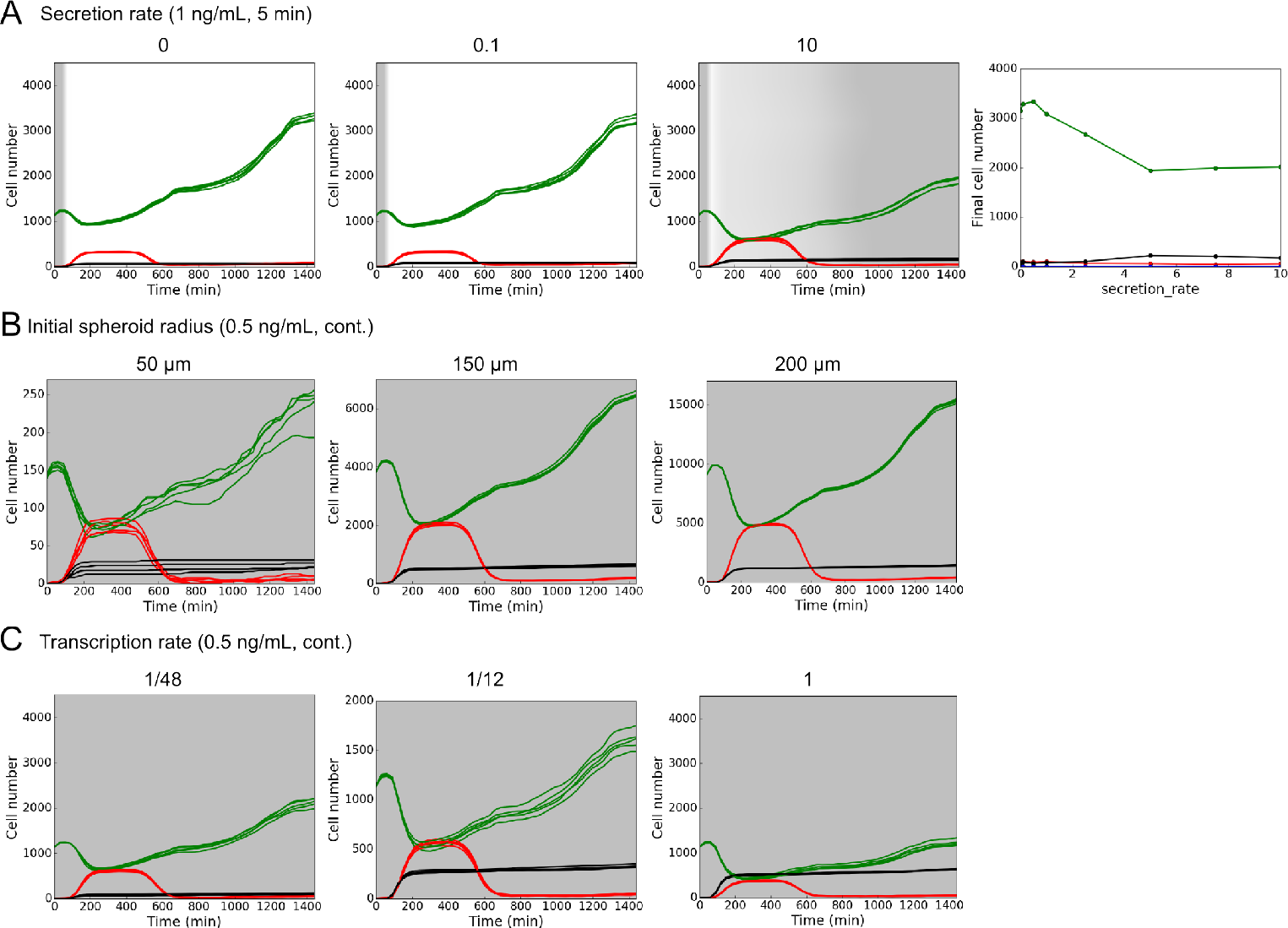
Effects of simulation parameters. A: Variation of secretion rate value. Time
evolution of the number of cells in each fate (left) for a secretion rate of: 0 (1st graphic), 0.1 (2nd) and 10 (3rd). TNF was injected at the beginning of the simulation during 5 min at a dose of 1 ng/mL. Final number of cells in each fate according to the secretion rate used for individual simulations (right). Simulation time is 12h, initial disk radius is 400 *μ*m, which accounts for roughly 1000 cells. B: Variation of the initial spheroid radius. Time evolution of the number of cells in each cell fate for an initial population radius of 50 (left), 150 (middle) and 200 (right) *μ*m. TNF is injected continuously during the simulations at a concentration of 0.5 ng/mL. Simulation time is 12h, note that Y-axis is different for the three graphs. C: Variation of the transcription rate used in the Boolean network transitions. Time evolution of the number of cell in each fate for a transcription rate of 1/48 (left), 1/12 (middle) and 1 (right). Other transitions rate are 1. Simulation time is 12h, initial disk radius is 400 μm, which accounts for roughly 1000 cells.A-C: Colour code: green, Proliferative cells; red, cells committed to Apoptosis; black, cells committed to NonACD. Grey shading indicates TNF presence. Except in panel A, right graph, 5 simulations are presented for each condition.

**S3 Fig.**
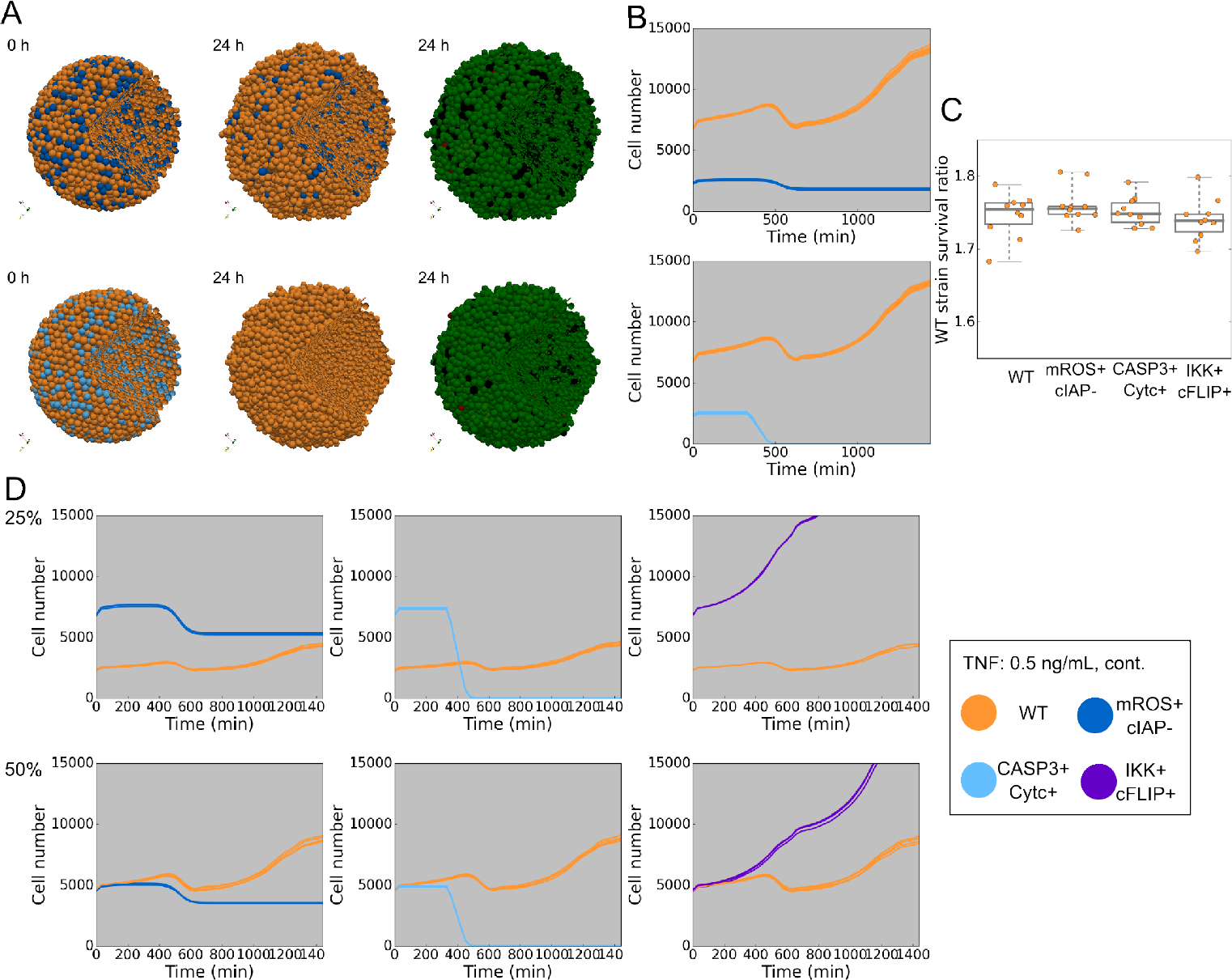
Genetically heterogeneous population under TNF treatment. Simulations of heterogeneous population composed of 75% of WT cells (orange) and 25% of mROS+ and cIAP- mutated cells (blue) or with CASP3+ and Cytc+ mutated cells (light blue). A: Snapshots of a simulation for each case at initial and final time (24 h), with cells coloured by cell type (left and middle) or by cell fate commitment (right). B: Time evolution of the number of cells of each strain (WT and mutated) for 10 simulations. C: Ratio of final number of surviving cells against initial number of cells for each cell line (WT or mutated). D: Time evolution of the number of cells in each strain (WT and mutated) for 10 simulations for the 3 different mutants with different initial proportion of WT cells compared to the total population: 25% (top) and 50% (bottom). A-C: Cell fate commitment colours: green, Proliferative cells; red, Apoptosis; black, NonACD. Grey shading indicates presence of TNF in continuous injection at 0.5 ng/mL. Initial spheroid radius is 200 *μ*m, which accounts for roughly 9000 cells, + stands for over-expression and - stands for knock-out.

**S4 Fig.**
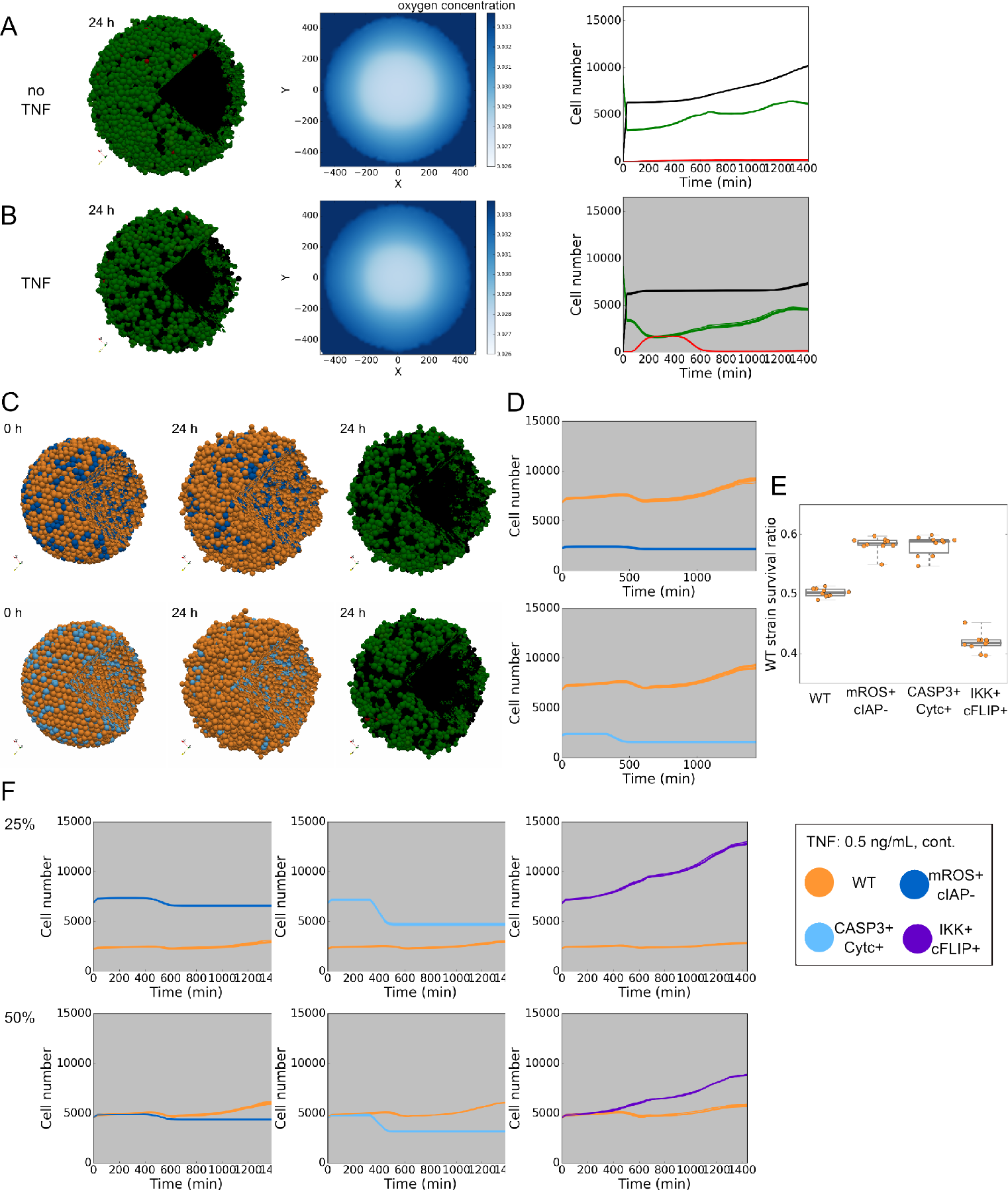
Genetically heterogeneous population under TNF treatment and oxygen-limited regime. A: Simulation of WT-only population without TNF. Snapshots of a simulation (left) and oxygen levels (middle) at 24 h. Time evolution of the number of cells in each cell fate (right). B: Simulation of homogeneous population for a low-dose injection of TNF. Snapshots of a simulation (left) and oxygen levels (middle) at 24 h. Time evolution of the number of cells in each cell fate (right). C-E: Simulations of heterogeneous population and TNF treatment composed of 75% of WT cells (orange) and 25% of mROS+ and cIAP- mutated cells (blue) or with CASP3+ and Cytc+ mutated cells (light blue). C: Snapshots of a simulation for each case at initial and final time (24 h), with cells coloured by cell type (left and middle) or by cell fate commitment (right). D: Time evolution of the number of cells in each strain (WT and mutated) for 10 simulations. E: Ratio of final number of surviving cells against initial number of cells for each cell line (WT or mutated). F: Time evolution of the number of cells in each strain (WT and mutated) for 10 simulations for the 3 different mutants with different initial proportion of WT cells compared to the total population: 25% (top) and 50% (bottom). A-F: Oxygen levels, represented from dark blue (injected level) to white (lowest level), are measured in the z=0 plane of the simulated space. Cell fate commitment colours: green, Proliferative cells; red, Apoptosis; black, NonACD. Grey level background in graphs indicates presence of TNF in continuous injection at 0.5 ng/mL. All simulations are in oxygen-limited regime, initial spheroid radius is 200 *μ*m, which accounts for roughly 9000 cells, + stands for over-expression and - stands for knock-out.

**S1 File. Supplementary information of PhysiBoSS implementation.** Additional details of PhysiBoSS usage, its implementation, and how the interface between the agent-based part and Boolean network part is handled are presented in this file.

**S2 File. Supplementary information of the cell fate simulations.** Additional details on cell fate study part. Description of the Boolean network used, short description of the pipeline used to identify interesting mutants, description of how TNF injection are simulated and parameters used in the simulations are presented in this file.

**S1 Table.** **Simulation run time.** Representative examples of simulation run time necessary for different kind of simulations. Simulations were run on one node of a Linux cluster (with 16 openMP threads).

| Cell number (init-final) | Simulated time | Remark | Run time |
| --- | --- | --- | --- |
| 1022 - 1969 | 720 min | no TNF | 2 min |
| 1024 - 1443 | 720 min | cont. TNF | 1.5 min |
| 1131 - 4772 | 1440 min | no TNF | 10.5 min |
| 1134 - 2211 | 1440 min | cont. TNF | 25.75 min |
| 1136 - 3544 | 1440 min | one pulse injection | 9 min |
| 3843 - 7284 | 1440 min | cont. TNF | 42 min |
| 16297 - 23227 | 1440 min | 7192 passive cells | 45.5 min |

